# The flexible N-terminus of BchL protects its [4Fe-4S] cluster in oxygenic environments and autoinhibits activity

**DOI:** 10.1101/840439

**Authors:** Elliot Corless, Syed Muhammad Saad Imran, Maxwell B. Watkins, Sofia Origanti, John-Paul Bacik, Robert Kitelinger, Mark Soffe, Karamatullah Danyal, Lance C. Seefeldt, Brian Bennett, Nozomi Ando, Edwin Antony

## Abstract

The dark-operative protochlorophyllide oxidoreductase (DPOR) enzyme contains two [4Fe-4S]- containing component proteins (BchL and BchNB) that assemble in an ATP-dependent fashion to coordinate electron transfer and reduction of protochlorophyllide to chlorophyllide. Photosynthesis generates an oxygenic environment that is non-optimal for [Fe-S] clusters and we here present an elegant evolutionarily conserved mechanism in BchL to protect its [4Fe-4S] cluster. We present a crystal structure of BchL in the nucleotide-free form with an ordered N-terminus that shields the [4Fe-4S] cluster at the docking interface between BchL and BchNB. Amino acid substitutions that perturb the shielding of the [4Fe-4S] cluster produce an unstable, but hyper-active enzyme complex, suggesting a role for the N-terminus in both auto-inhibition and enzyme stability. Upon ATP binding, a patch of amino acids, Asp-Phe-Asp (‘DFD patch’), situated at the mouth of the BchL ATP-binding pocket promotes inter-subunit cross stabilization of the two subunits and relieves the auto-inhibition by the N-terminus. A linked BchL dimer with one defective ATP-binding site does not support substrate reduction, illustrating that nucleotide binding to both subunits is a prerequisite for the inter-subunit cross stabilization. We propose that ATP-binding produces a conformational compaction of the BchL homodimer leading to a release of the flexible N-terminus from blocking the [4Fe-4S] cluster and promotes complex formation with BchNB to drive electron transfer. The auto-inhibitive feature and release mechanism appear unique to DPOR and is not found in the structurally similar nitrogenase.

## Introduction

Photosynthetic organisms utilize chlorophyll or bacteriochlorophyll to capture light for their energy requirements. The multi-step enzymatic biosynthesis of both these compounds are similar in the cell except for the penultimate reduction of protochlorophyllide (Pchlide) to produce chlorophyllide (Chlide).^1,2^ Angiosperms use a light-dependent protochlorophyllide oxidoreductase to catalyze the reduction, whereas gymnosperms, cyanobacteria, algae, bryophytes and pteridophytes possess a light-independent enzyme called dark-operative protochlorophyllide oxidoreductase (DPOR; Fig. 1a).^3^ Photosynthetic bacteria that are anoxygenic, such as *Rhodobacter capsulatus*, rely exclusively on the activity of DPOR for synthesis of bacteriochlorophyll.^3^ DPOR catalyzes the stereospecific reduction of the C17 = C18 double bond of Pchlide to form Chlide (Fig. 1b). This reduction forms the conjugated π-system in the chlorin structure of chlorophyll-a which leads to a parental shift in the spectral properties required for photosynthesis.^4,5^

**Figure 1.**
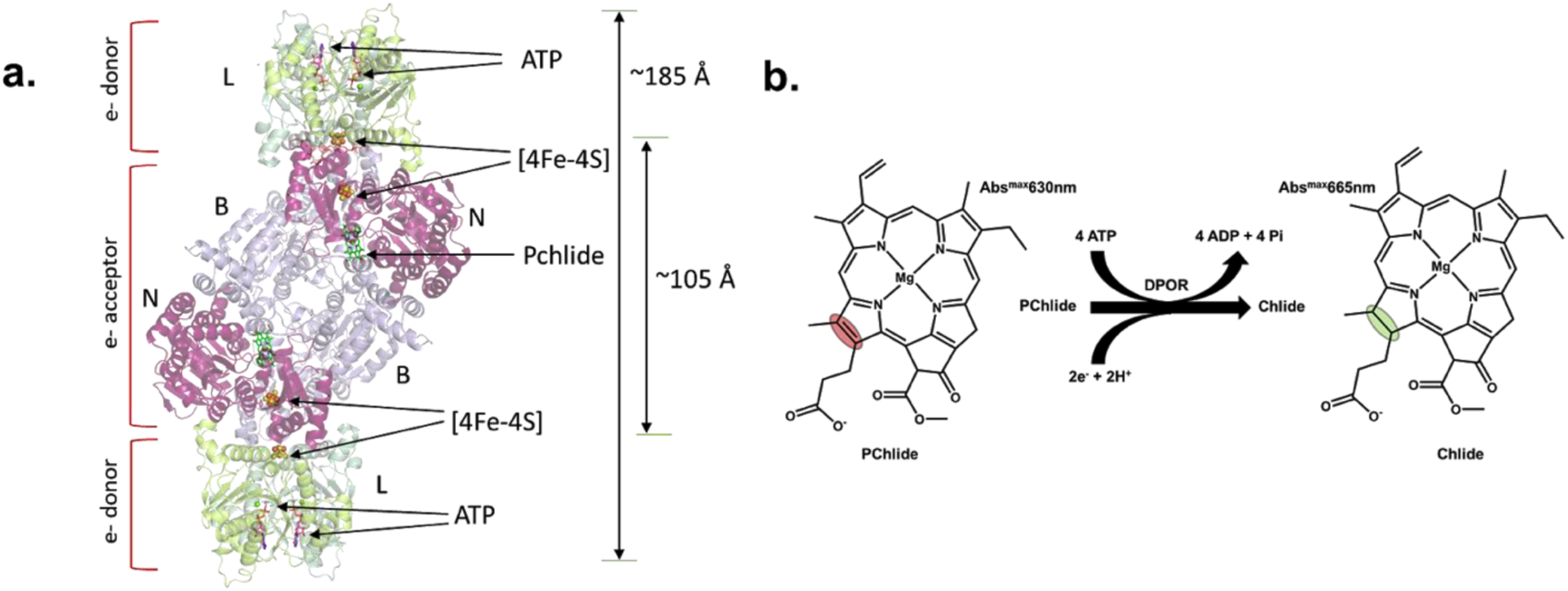
Structure and substrate reduction mechanism of DPOR. (**a**) Crystal structure of the complete, ADP-AlF_3_-stabilized, DPOR complex (PDB:2ynm). BchL subunits are colored green, BchN is colored purple, and BchB is violet. The four [4Fe-4S] clusters are shown as spheres, and ADP-AlF_3_ and Pchlide are shown as sticks. (**b**) Schematic of Pchlide reduction to Chlide by DPOR. Two cycles of electron transfer from BchL to BchNB are required for the reduction of the C17=C18 double bond (marked by the colored ovals).

DPOR is structurally homologous to nitrogenase, the enzyme responsible for reducing di-nitrogen to ammonia. DPOR is composed of two components: a homodimeric L-protein (BchL) and an heterotetrameric NB-protein (BchNB).^5,6^ BchL serves as the ATP-dependent electron donor, and BchNB is the electron acceptor containing the active site for Pchlide binding and reduction (Fig. 1a). A multi-step reaction cycle has been proposed for DPOR function with the following overall reaction stoichiometry (Fig. 1b)^4^:

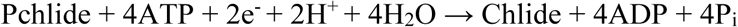

Give the structural similarity to nitrogenase, ATP binding to BchL is thought to promote its transient association with BchNB followed by a single electron transfer (ET) to Pchlide. ATP hydrolysis drives the dissociation of the protein complex. Two such rounds of ATP-dependent ET are necessary for Pchlide reduction, and the minimum stoichiometry of ATP/molecule of Chlide formed has been determined to be 4.^7^ However, the details of how ATP binding promotes complex formation between BchL and BchNB are poorly resolved.

High-resolution crystal structures of the BchL dimer complexed with ADP (PDB:3FWY)^8^, and stabilized in a higher-order complex with the BchNB tetramer by ADP-AlF_3_ (PDB:2YNM; Fig. 1a)^9^ provide several key molecular insights: BchL contains one [4Fe-4S] cluster ligated by 2 conserved cysteine residues from each subunit of the dimer. The subunits of BchL each have an active site for ATP, including conserved binding (Walker A) and hydrolysis (Walker B) motifs. Each half of the BchNB tetramer contains a substrate (Pchlide) binding site and one [4Fe-4S] cluster which functions as the electron acceptor from BchL (Fig. 1a). This cluster is ligated by 3 cysteine residues from BchN and one uncommon aspartic acid ligand from BchB. BchL sits across the top of BchNB, placing their metal clusters in relatively close proximity (∼ 16 Å; Fig. 1a)^9^. Thus, ATP binding is hypothesized to drive formation of the complex. ATP hydrolysis has been proposed to drive electron transfer. In the homologous nitrogenase system, ATP hydrolysis occurs post-ET suggesting that hydrolysis likely drives complex dissociation post-ET.^10^ Given the structural similarities between DPOR and nitrogenase, we hypothesize ATP hydrolysis is also likely to promote complex dissociation in DPOR. Here, we address three key questions about ATP usage by BchL: How does binding of 2 ATP molecules collectively transmit information from the ATP binding sites to the [4Fe-4S] cluster of BchL along with the interface where it complexes with BchNB? What role does ATP play in electron transfer? Are both ATP binding events necessary?

We present a crystal structure of *Rhodobacter sphaeroides* BchL in the nucleotide-free state. This structure reveals novel electron density for a flexible N-terminal region that is bound across the face of the BchL [4Fe-4S] cluster, suggesting a potential regulatory role. We show that amino acid substitutions within this flexible N-terminal region enhance the kinetics of substrate reduction, pointing to a functionally suppressive role for this interaction. Additionally, inter-subunit contacts between BchL and the bound-ATP are critical for substrate reduction activity. Finally, we show that ATP binding to both subunits is required to promote conformational changes requisite for reduction of Pchlide to Chlide. We propose a model where ATP-driven cross stabilization of the homodimer promotes the release of the flexible N-terminus and drives formation of the DPOR complex towards ET and substrate reduction.

## Results

### Crystal structure of nucleotide-free BchL suggests regulation through a redox switch and a flexible N-terminal region

BchL was crystallized anaerobically in the absence of nucleotides, and a crystal structure was determined to a resolution of 2.6 Å (Supplemental Table 1). The asymmetric unit contains four BchL chains: chains A and B form one BchL dimer, and chains C and D comprise the other (Supplemental Fig. 1a,b). The overall structure of BchL is similar to previously published crystal structures (Fig. 2a-c), but in this nucleotide-free state, the top face of the dimer is more open compared to ADP-bound BchL^8^ (Fig. 2b) and ADP-AlF_3_-NB-bound BchL^9^ (Fig. 2c). Unexpectedly, although the N-terminus is disordered and not observed in all previous BchL structures (residues 1-29 in 3FWY and residues 1-27 in 2YNM from *P. marinus*), we observe clear electron density at the N-terminus of chain C in our nucleotide-free structure, which we were able to model as residues 16 to 29 (Supplemental Fig. 1b). Interestingly, the flexible N-terminal region is observed bound across the [4Fe-4S] cluster in our structure (Fig. 2a) covering a surface that is normally used to interface with BchNB (Fig. 2c). In the chain C/D dimer, Asp23 of the N-terminal region interacts with Gln168, Gly16 forms an H-bond to Cys126 (an [4Fe-4S] cluster ligating residue) of the opposing chain, while Ser17 interacts with Gly161 of chain C via the backbone of G125 in chain D (Fig. 2d, Supplemental Fig. 1c). Based on the structure of the ADP-AlF_3_-stabilized DPOR complex (PDB: 2NYM)^9^, the residues corresponding to Cys126, Gly161, and Gln168 in BchL (*R. sphaeroides* numbering) are three of the twelve residues that interact with BchNB during the formation of the active complex, as predicted by PDBePISA interface analysis^11^ (Fig. 2c, purple surface). Notably, the flexible N-terminal protective region (residues 1-29) is only conserved among DPOR BchL-proteins and is not observed in other homologous proteins such as nitrogenase and the BchX protein of chlorophyllide oxidoreductase (Supplemental Fig. 2). The position and interactions of the N-terminal residues in our nucleotide-free structure suggests a possible auto-inhibitory role by forming a barrier to docking and shielding the [4Fe-4S] cluster of BchL.

**Figure 2.**
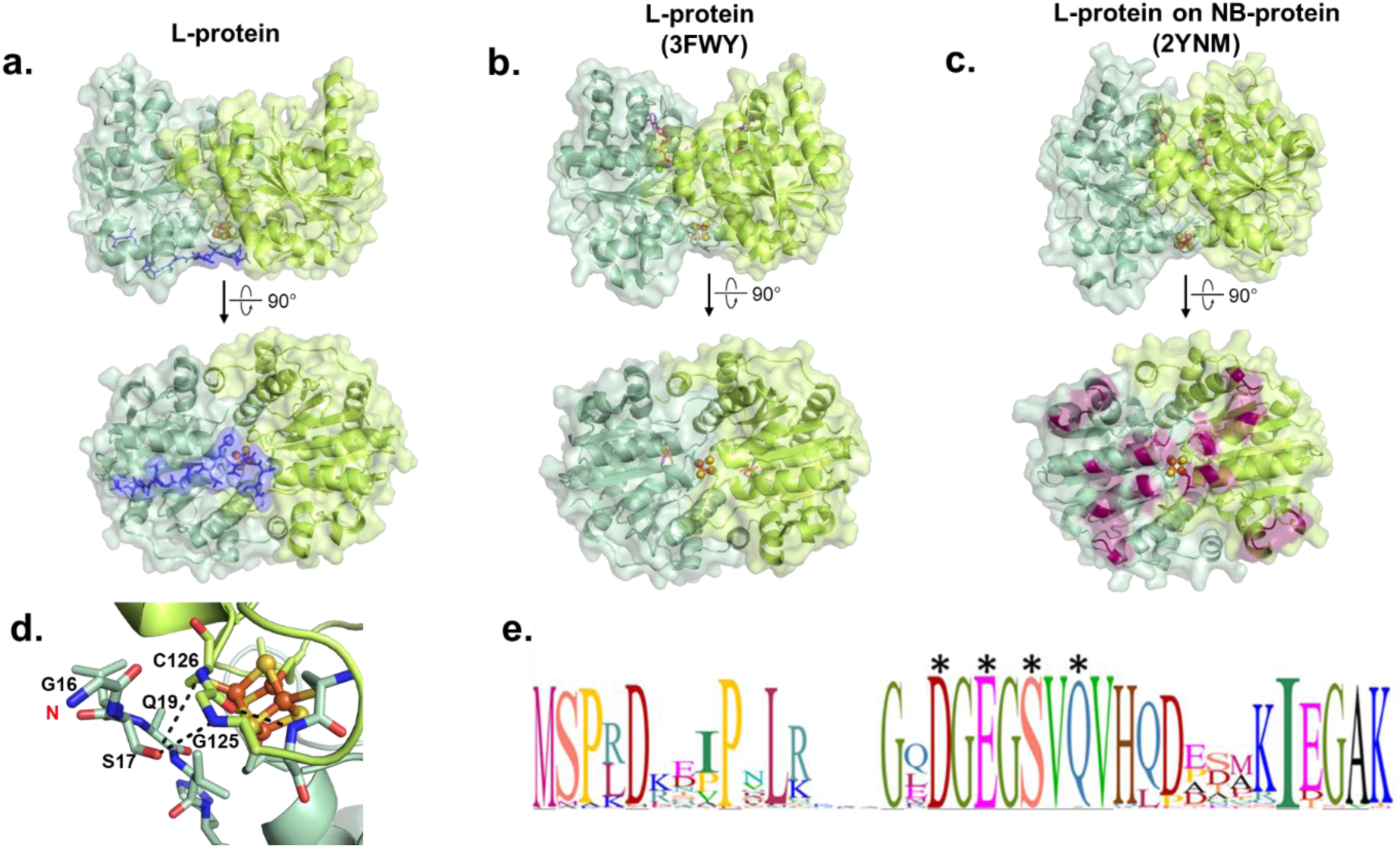
Crystal structure of nucleotide-free BchL reveals a flexible N-terminal region capping the [4Fe-4S] cluster. Side views (top row) and bottom views (bottom row) of the BchL structure (**a**) in the absence of nucleotides, (**b**) with ADP bound (PDB: 2YNM), and (**c**) in complex with BchNB (PDB: 3FWY). A slight compaction upon the addition of ADP, and a further compaction when in complex with BchNB, can clearly be seen by comparing the side views. The flexible N-terminal region resolved in the nucleotide-free structure (residues 16-29, colored blue in panel a) clearly covers the [4Fe-4S] cluster in addition to blocking or directly interacting with several residues predicted to directly interact with BchNB (highlighted in dark purple in panel c). **(d)** Residues in the flexible N-terminus of chain C interact with important residues near the [4Fe-4S] cluster. The highly conserved Ser17 forms hydrogen bonds with both Cys126 and Gly125 on chain C, the former of which directly interacts with BchNB in the 2YNM structure and the latter additionally interacts with Gly161 on chain D, another predicted BchL-BchNB interface residue. **(e)** Sequence logo of the N-terminus of BchL generated from alignment of n=89 species. Letter height corresponds to the degree of sequence conservation, and the residues mutated in this study are labeled.

### Amino acid substitutions in the flexible N-terminal region of BchL increases the rate of substrate reduction

To test whether the flexible N-terminal region of BchL plays an auto-inhibitory role, based on the contacts observed in the crystal structure (Fig. 2d) and sequence alignments (Fig. 2e), we generated a singly mutated construct (BchL^S17A^) where Ser17 was mutated to Ala, and a quadruple mutated construct (BchL^4A^) where Asp13, Glu15, Ser17 and Gln19 were all substituted with Ala. Though density for residues 1-15 is not seen in any BchL structure, Ser17 and Gln19 are observed in our structure positioned across the [4Fe-4S] cluster (Supplemental Fig. 1b). Clarified cell lysates containing BchL^S17A^ was mixed with Pchlide and BchNB and found to possess reductase activity after purification (Fig. 3b). However, amino acids substitutions in this N-terminal region affected protein stability: Both BchL^S17A^ and BchL^4A^ are soluble to a lesser degree compared to the wild-type protein and formed cloudy precipitates within 30 minutes of initiating reduction reactions. Substrate reducing activity with purified protein was assayed by mixing purified BchL, BchL^S17A^, or BchL^4A^ (4 μM) with BchNB (1 μM) and Pchlide (35 μM) and spectroscopically monitoring the reaction over time in the absence or presence of ATP (3 mM). Pchlide and Chlide have characteristic absorbance maxima at 630 nm and 680 nm, respectively, in aqueous solution. Formation of Chlide was observed as an increase in absorbance at 680 nm in the presence of ATP (Fig. 3b). Both BchL^S17A^ and BchL^4A^ are active for substrate reduction and show Chlide formation rates ∼2-2.5 fold that of wild type BchL (k_obs_= 0.044 ± 0.014, 0.084 ± 0.019, and 0.087 ± 0.016 µM/min for BchL, BchL^S17A^ and BchL^4A^, respectively; Fig. 3c). We speculate that actual differences in activity are likely much larger as the effective concentrations of the mutant BchL proteins are likely lower than calculated during the experiment due to protein instability. We also generated a truncated version of BchL missing the first 27 amino acids (BchL^NΔ27^). We were successfully able to overexpress BchL^NΔ27^, however it was poorly soluble, and we were unable to obtain sufficiently pure protein for biochemical studies. These difficulties provide additional evidence that the flexible N-terminus might play an important role in BchL function and stability. We also recently showed that the presence of a functional Suf operon in *E. coli* cells leads to a 3-fold enhancement of BchL overproduction suggesting that the flexible N-terminus might be modulated by specific cellular mechanisms.^12^

**Figure 3.**
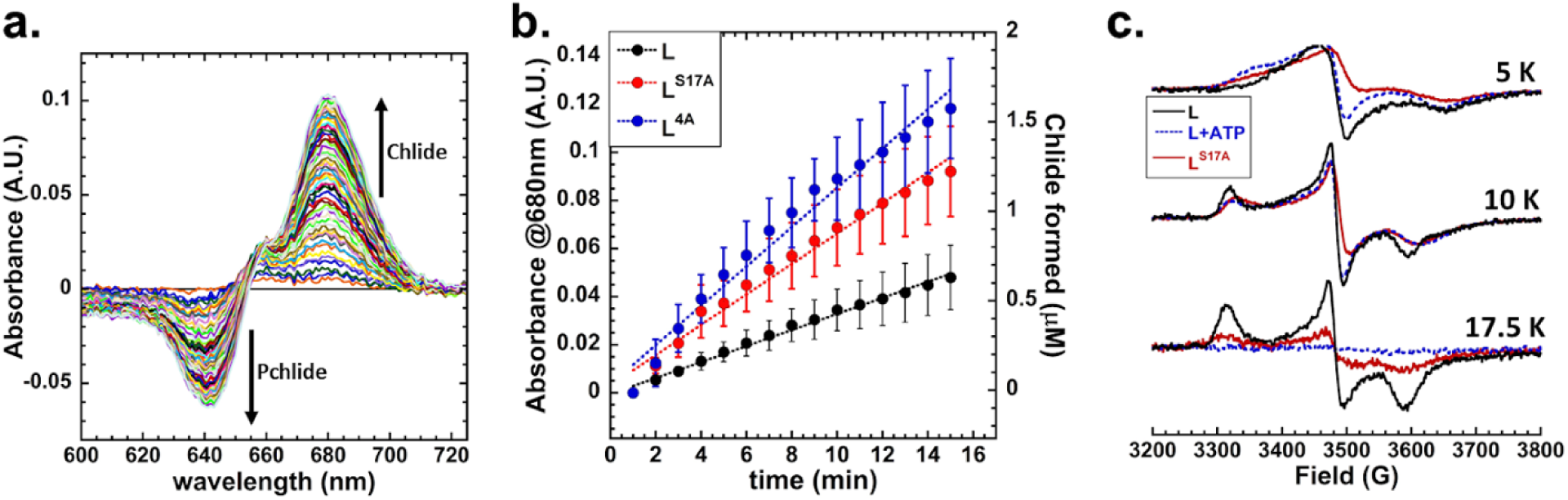
The flexible N-terminal region is auto-inhibitory to BchL function. **(a)** Representative traces of wild-type reaction in aqueous solution. Arrows represent the trend in absorbance associated with substrate consumption and product formation. **(b)** Absorbance plot at 680nm (A680) of the *in vitro* reaction comparing reduction rates as a change in A_680_ (left axis) and apparent Chlide formation rates (right axis) of BchL (black circles, *k_obs_*=0.044 ± 0.014 µM.min^-1^) BchL^S17A^ (red circles, *k_obs_* =0.084 ± 0.019 µM.min^-1^), and BchL^4A^ (blue circles, *k_obs_* =0.087 ± 0.016 µM.min^-1^) **(c)** EPR spectra comparing BchL (black solid lines) BchL incubated with excess ATP (blue dotted lines), and BchL^S17A^ (Red solid lines) at 5 K, 10 K and 17.5 K where indicated.

### ATP-binding causes changes in the local environment of the BchL [4Fe-4S] cluster affecting their EPR spectral line-shape and intensities

Since the flexible N-terminal region binds to the BchNB interaction interface within BchL, we hypothesized that ATP binding could promote conformational changes in BchL to relieve autoinhibition. We thus used electron paramagnetic resonance (EPR) to probe local changes in the [4Fe-4S] cluster environment, which includes the binding site for the N-terminus. Interpretation of the g-tensors of [4Fe-4S] clusters in direct structural terms is rarely possible,^13^ but EPR can nevertheless provide information about relative conformational changes around the cluster. Previous studies have described the [4Fe-4S] cluster of BchL as an axial species,^5,6,14^ while others reported a rhombic species.^15,16^ Here, EPR at different temperatures indicated that two distinct EPR signals were exhibited by the [4Fe-4S] cluster (Supplemental Fig. 3a). At 5 K, a signal termed FeS^A^ was observed that appears axial but was best simulated with rhombic *g*-values of 2.00, 1.94, and 1.85; both *g*_1_ and *g*_3_ are atypically low for a prototypical Cys_4_-ligated [4Fe-4S] cluster, and the associated resonances exhibit large line widths (Supplemental Fig. 3a). This signal was very fast-relaxing and was no longer detectable at 17.5 K. At this higher temperature (17.5 K), a more typical rhombic signal, FeS^B^, with *g*_1,2,3_ = 2.04, 1.94, and 1.89 was observed, and at an intermediate temperature (10 K) the observed signal was well replicated by a 40%:60% mixture of the simulations of FeS^A^ and FeS^B^, respectively. As expected for a [4Fe-4S] cluster, the EPR signals were fast-relaxing and were not detectable at 30 K or higher temperature. BchL exhibited analogous signals to FeS^A^ and FeS^B^ when incubated with BchNB and ADP (Supplemental Figs. 3b and 3c) although, notably, both ADP and BchNB served to increase the relaxation rate of the FeS^B^ EPR signal, suggesting a more efficient coupling of the cluster to the lattice via increased strain energy. In addition, with ADP, the proportion of the FeS^A^ species was diminished by a factor of two. Based on the structural data for BchL and the relaxation properties of the EPR signals, we propose the FeS^A^ species as having a ‘cap’ across the cluster, formed by the flexible N-terminus, whereas the FeS^B^ species is uncapped; in solution, these two species are likely in dynamic equilibrium. Upon the addition of ATP, the relaxation rate of the FeS^B^ species was further enhanced and was undetectable at 17.5 K (Supplemental Fig. 3c), while the FeS^A^ signal exhibited rapid-passage distortion at 5 K, indicating a diminution of the relaxation rate for that species. These data suggest that ATP binding increases the conformational strain of uncapped FeS^B^ and somewhat inhibits the strong interaction of the cap with the cluster in FeS^A^. The FeS^A^ EPR signals from the BchL^S17A^ variant (Supplemental Fig. 3C) exhibited strong rapid-passage distortion at 5 K and overall reduced signal intensities over the 5 - 17.5 K temperature range; the addition of ATP restored the intensity of the FeS^B^ signal somewhat, suggesting that relaxation properties were responsible for this phenomenon and that, therefore, the interaction of the cap region with the cluster is altered in BchL^S17A^. The conformational changes around the [4Fe-4S] cluster in BchL upon ATP binding appear similar to the BchL^S17A^ protein in the absence of ATP (Fig. 3c). Based on these results, we propose that the loss of auto-inhibition drives higher overall substrate reduction activity in BchL^S17A^.

### A DFD amino-acid patch promotes inter-subunit cross-stabilization upon ATP binding

The ATP-binding sites in BchL are situated away from the BchL:BchNB interaction interface (Fig. 1a, Fig. 2a-c). Thus, we hypothesized that other regions between these two sites might function as a conduit for communication and be an important contributor to potential conformational changes and promote rearrangement of the flexible N-terminus. Comparison of the nucleotide-free (Fig. 4a), ADP-bound (Fig. 4b)^8^ and ADP-AlF_3_-NB-bound (Fig. 4c)^9^ BchL crystal structures reveal a ‘DFD patch’ composed of amino acid residues Asp180, Phe181, Asp182 that undergo rearrangements and contact the hydroxyl groups of the sugar moiety of the bound nucleotide. Interestingly, the contacts only appear in the ADP-AlF_3_-L-NB DPOR structure (Fig. 4c), and thus the DFD patch appears poised to be important for coordinating ATP-dependent conformational changes during complex formation and substrate reduction. D180 and D182 from one subunit form a network of interactions in *trans* with the sugar moiety of ATP bound to the neighboring subunit along with R244 in the nucleotide-bound subunit (Fig. 4c). We have termed this series of interactions “inter-subunit cross stabilization of ATP”. These interactions appear to be stabilized in the transition state when BchL forms a complex with BchNB. One possibility is that ATP-binding drives inter-subunit cross stabilization and would provide the necessary conformational stability required to pry away the flexible N-terminal tail from binding across the [4Fe-4S] cluster, thus relieving inhibition. If this were the case, mutations in the DFD patch would not affect ATP binding, but would perturb substrate reduction.

**Figure 4.**
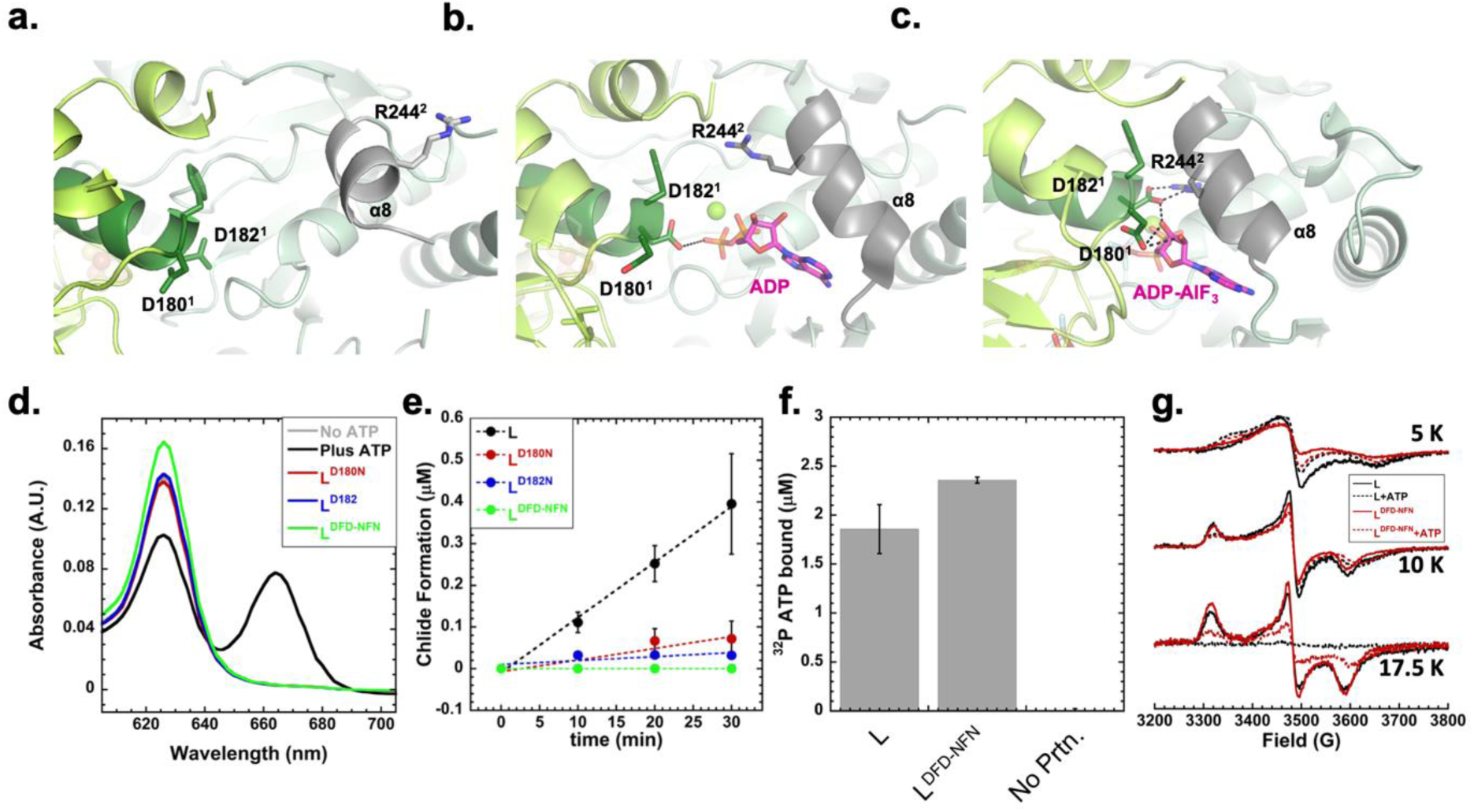
Interactions between ATP and a DFD patch promotes inter-subunit cross stabilization and conformational changes in BchL. Views of the DFD patch (highlighted in green) and associated interactions in three distinct conformations of BchL. **(a)** In the absence of nucleotides, residues D180 and D182 on chain C form no notable interactions and are a large distance away from residue R244 on the opposing chain D. **(b)** With ADP bound, the α8-helix, highlighted in dark grey. undergoes considerable motion, bringing R244 much closer to the DFD patch. Additionally, D182 now interacts with the bound ADP. **(c)** When bound to ADP-AlF_3_ and in complex with BchNB, D180 and D182 have extensive interactions with both the bound nucleotide as well inter-subunit interactions with R244. **(d)** Pchlide reduction activity of BchL and mutant proteins. Chlide formation is monitored as a peak between 660-670 nm and occurs only in the presence of ATP. BchL^D180N^, BchL^D182N^, and BchL^DFD-NFN^, are all defective for substrate reduction. **(e)** Time course of Chlide formation for reactions containing BchNB, ATP and Pchlide in the presence of various BchL constructs. BchL reduces Pchlide to Chlide (*k_obs_* s=0.0126 ± 0065 µM.min^-1^), and no appreciable Chlide formation is observed for BchL^D182N^ or BchL^DFD-NFN^. Severely impaired but detectable reduction activity is observed for BchL^D180N^ (*k_obs_* =0.0028 ± 0.0042 µM.min^-1^). **(f)** Nitrocellulose filter binding analysis of ATP binding to BchL shows ∼2 ATP bound per BchL dimer to both BchL and BchL^DFD-NFN^, and no non-specific binding to the membranes is observed in the absence of BchL in the reaction. **(g)** EPR spectra comparing wt BchL (black solid lines), BchL incubated with excess ATP (black dotted lines), BchL^DFD-NFN^ (Red solid lines), and BchL^DFD-NFN^ incubated with excess ATP (red dotted lines) at 5 K, 10K, and 17.5K as denoted.

To test this hypothesis, and the role of the DFD patch, we generated three BchL variants carrying amino acid substitutions wherein Asp180 (BchL^D180N^), Asp182 (BchL^D182N^), or both, (BchL^DFD-NFN^) were substituted to Asn. Phe181 does not make contacts with the sugar and thus was not perturbed. BchL^D180N^ was poorly active for Pchlide reduction (Fig. 4d, e) whereas BchL^D182N^ and BchL^DFD-NFN^ were inactive (Fig. 4d, e), suggesting that both residues are important for function. Next, to precisely understand why mutations in the DFD patch affected substrate reduction, we analyzed the nucleotide binding, and EPR spectral properties of BchL^DFD-NFN^. Nucleotide binding was measured by capturing the BchL:ATP complex using radiolabeled α^32^P-ATP in a nitrocellulose filter binding assay. BchL binds to the nitrocellulose filter and ATP bound to the protein is retained on the membrane, while unbound ATP flows through the filter. Both BchL and BchL^DFD-DFN^ are capable of binding to ATP, and no ATP is retained on the membrane when no protein is present in the reaction (Fig. 4f). This finding is consistent with the ADP-bound crystal structure where no contacts between the DFD patch and nucleotide are observed (Fig. 4b). Thus, the loss in substrate reduction activity in the BchL^DFD-NFN^ protein occurs post ATP binding, and possibly due to a loss of the promotion of conformational changes necessary for multiple rounds of complex formation with BchNB.

EPR of BchL^DFD-NFN^ at 5 K (Supplemental Figure 3e and Fig. 4g) again revealed an FeS^A^ signal, though the rapid-passage distortions indicated that ATP binding enhanced relaxation in BchL^DFD-NFN^ whereas ATP inhibits relaxation in native BchL. The overall relaxation rates for FeS^A^ in native BchL and BchL^DFD-NFN^ can be summarized as BchL > BchL-ATP ≈ BchL^DFD-NFN^-ATP > BchL^DFD-NFN^. At 10 K, the EPR spectra of both native BchL and BchL^DFD-NFN^ are almost indistinguishable and consist of 60% FeS^B^, whereas the ATP complexes of both exhibit EPR signals indistinguishable from each other but containing only 35% FeS^B^ and 65 % of the relaxation-inhibited FeS^A^ (Supplemental Figure 5e). The spectra of both native BchL and BchL^DFD-NFN^ at 17.5 K are again indistinguishable and are due to FeS^B^ alone (Supplemental Figure 5e). In both cases, the signals are of diminished intensity upon addition of ATP due to enhanced relaxation, to about 25% in the case of BchL^DFD-NFN^-ATP and almost completely extinguished in native BchL-ATP. It is not clear whether the residual FeS^B^ signal from BchL^DFD-NFN^-ATP at 17.5 K is due to slower relaxation than in native BchL-ATP or less than stoichiometric binding of ATP. So, overall, relaxation of FeS^A^ is markedly inhibited in BchL^DFD-NFN^, indicating poorer coupling to a strained lattice in the capped species, while the cluster environments in the uncapped FeS^B^ species of native BchL and BchL^DFD-NFN^ are indistinguishable by EPR (Fig. 4g); and the binding of ATP to both species provides very similar cluster environments for both FeS^A^ and FeS^B^. Thus, ATP binding to BchL^DFD-NFN^ does not elicit the complete portfolio of conformational changes required for substrate reduction.

### Binding of ATP to both subunits in the BchL dimer are required to generate a concerted motion to promote inter-subunit cross stabilization and drive substrate reduction

Each BchL homodimer consists of two sites for ATP binding (Fig. 5a), and the two subunits are covalently tethered by a single [4Fe-4S] cluster (Fig. 2a-c). In the homologous Fe protein of nitrogenase, the sites have been shown to bind to nucleotide with differing affinities.^17^ A crystal structure of the Fe-protein with two different nucleotides occupying the dimer has also been solved suggesting that the two ATP sites could play distinct roles in substrate reduction^18^ (Supplemental Fig. 3c). To test the functional role of the two ATP-binding sites in BchL, we generated a covalently linked version of BchL by expressing the two subunits as a single polypeptide. The “linked” construct included the same N-terminal 6x-poly histidine tag followed by a TEV protease site identical to all other L-protein constructs. Two BchL subunits were tethered by a flexible linker that connected the C-terminal end of the first monomer to the N-terminal end of the other (Fig. 5b). Covalent linkage of multimers can interfere with activity due to a variety of effects including conformational strain, non-specific interactions due to the linker, introduction of non-native secondary structures, or undefined effects. The linker was therefore constructed to carry TEV protease recognition sites bookending the linker region enabling proteolytic removal if the intact linker interfered with activity for any reason. (Fig. 5b). We found that the length of the linker is a key determinant of protein activity. Linkers shorter than 15 amino acids are defective for substrate reduction, and optimal activity is obtained when linker lengths are longer than 20 amino acids (Supplemental Fig. 4c, d). The optimized linked-BchL behaved similarly to wild-type BchL during purification (Fig. 5c) and was stable and fully active for Pchlide reduction (Fig. 5d). Removal of the linker after protease cleavage also resulted in protein activity similar to the uncleaved and wild-type BchL proteins (Supplemental Fig. 4b). These results show that a linker of optimal length does not interfere with protein function. The linked L-protein appeared to have slightly faster activity. We believe that this is due to only one N-terminal tail being free to “cap” the [4Fe-4S] cluster, whilst the other is attached to the C-terminus of the first monomer and likely does not contribute to auto-inhibition of activity. This suggests that the 2 available N-terminal tails in the unlinked construct are dynamically interacting with the cluster.

**Figure 5.**
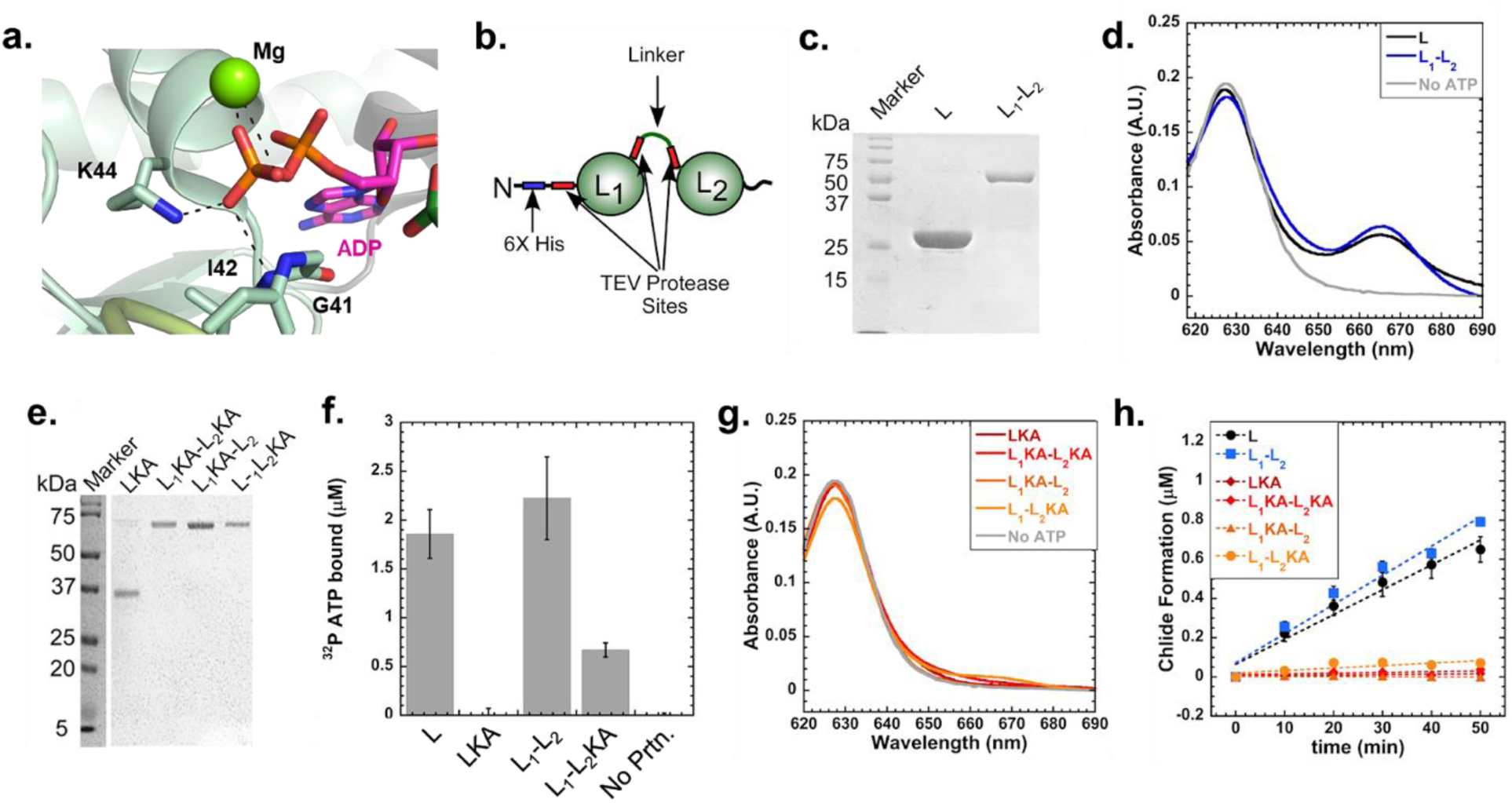
Both ATP binding sites are required for DPOR function. (**a**) Lys44 stabilizes nucleotide binding through interactions with the phosphate group of ADP (PDB: 3FWY). (**b**) Schematic of the linked-L-protein design and the positions of the tobacco etch virus (TEV) protease cleavage sites. (**c**) SDS-PAGE analysis of the purified BchL and linked-BchL-proteins. (**d**) Spectroscopic analysis of Pchlide reduction activity of BchL (L, black trace), linked-BchL-protein (L_1_-L_2_, blue trace), and no ATP negative control (No ATP, grey trace). Pchlide absorbance is observed at 625 nm and Chlide formation is monitored at 665 nm. (**e**) SDS-PAGE analysis of the purified BchL^K44A^(LKA) and versions of the single and double Lys44 to Ala substituted linked-BchL-proteins. (**f**) Nitrocellulose filter binding analysis of ^32^P-ATP binding by wildtype and Lys44 to Ala substituted BchL proteins. L and L_1_-L_2_ are capable of binding ATP, whereas the singly substituted L_1_KA-L_2_ is partially able to bind ATP. When both subunits are substituted with Lys44 to Ala, no ATP binding is observed. (**g**) Linked and unlinked BchL proteins carrying the K44A substitution are incapable of reducing Pchlide. (**h**) Kinetics of Pchlide reduction measured as a function of Chlide formation is shown. Land L_1_-L_2_ reduce Pchlide to Chlide (*k_obs_* = 0.01266 ± 0.007 µM.min^-1^ and 0.0148 ± 0.0022 µM.min^-1^, respectively.)

Next, to assess the contribution of the individual ATP sites, we generated an ATP-binding deficient mutation in one or both subunits in linked-BchL (Fig. 5e). Substitution of the conserved Lys44 with an Ala in the Walker A motif perturbs ATP binding (Fig. 5a). Using filter binding analysis, we determined the amount of ATP bound to the linked and unlinked versions of BchL. BchL (L) and linked-BchL with unaltered ATP binding sites (L_1_L_2_) bind ATP to similar extents (Fig. 5f). The K44A substitution in the Walker-A ATP binding pocket of BchL (L^KA^) and in both subunits of the linked-BchL (L_1_^KA^-L_2_^KA^) abolishes ATP binding as expected (Fig. 5f). When only one of the two ATP binding sites are mutated (L_1_^KA^-L_2_), partial ATP binding is observed (Fig. 5f). When ATP binding is perturbed in both sites or in just one site, a complete loss of Pchlide reduction activity is observed (Fig. 5g, h). These data suggest that both ATP molecules in the BchL dimer are required for substrate reduction. Based on these data, we propose that binding of both ATP molecules likely causes cooperative conformational changes in the two halves of the L-protein homodimer and drive inter-subunit cross stabilization.

## Discussion

Nitrogenase and nitrogenase-like enzymes such as DPOR and Chlorophyllide Oxidoreductase (COR) share structural similarity with respect to their electron donor and electron acceptor component proteins. These proteins catalyze multiple rounds of ET for substrate reduction, and transient association of the electron donor and electron acceptor is a prerequisite for each ET event. ATP binding to the electron donor is canonically assigned as the mechanistic trigger that promotes the assembly of the component proteins. However, the precise structural and functional principles underlying this ATP-driven process have largely remained unclear. The results presented in this study shed light on several ATP-binding driven changes in BchL that enable DPOR to function in an oxygenic environment.

In the nucleotide-free crystal structure, residues 16-29 are ordered in one of the four chains and suggest a novel regulatory role for the flexible N-terminus of BchL. Interestingly, the N-terminal tail binds across the [4Fe-4S] cluster and appears poised to block the docking surface for interaction with BchNB (Fig. 2a-c). Six residues of each BchL chain are thought to interact with BchNB to form the DPOR complex. In our BchL structure, the disordered N-terminus from one subunit forms an inhibitory barrier across the docking surface, and hydrogen bonding affects three of the 12 docking residues (Cys 126, Gly 161, and Gln 168), suggesting that the N-terminus forms a barrier to docking and ET to BchNB. Mutations that perturb specific interactions in this region enhance the substrate reduction activity, supporting an auto-inhibitory role for this region in DPOR function (Fig. 3). Although binding of both ADP and ATP can be thought to trigger conformational changes leading to the displacement of N-terminal residues, only ATP hydrolysis drives the dissociation of the BchL-NB complex to complete the catalytic cycle. Therefore, the conformational change in the N-terminal region could be yet another way of coupling ATP-binding to BchL with Pchlide reduction in BchNB.

A BLAST-P analysis of residues 1-29 yielded BchL protein hits spread across ∼150 species of bacteria. A nine-residue patch in the flexible N-terminus [DGEGSVQVH] (residues 13-21 in *R. capsulatus* BchL) is highly conserved, supporting functional significance (Fig. 2e). The flexible N-terminal region is unique to the DPOR system. The tail is not conserved in COR which catalyzes the subsequent reductive step in chlorophyll synthesis (Fig. 2e and Supplemental Fig. 2), and the entire N-terminal region does not exist in the Fe-protein of nitrogenase. The biological necessity for such a regulatory region remains to be established, though may exist as a mechanism to prevent off-target electron donation and the formation of free radicals in the cell. The observations that substitutions in this region affect BchL stability *in vitro* suggests a potential role in protecting the [4Fe-4S] cluster against oxidation and off-target electron donation.

ATP binding to both subunits of BchL promotes a network of interactions between a conserved DFD patch and the sugar moiety of the nucleotide. This generates an inter-subunit cross stabilization as the DFD patch from one subunit contacts the nucleotide bound to the other. Such an ATP-binding dependent conformational change could generate an upward compaction of BchL leading to the release of the auto-inhibitory N-terminus from the docking interface and promote complex formation with BchNB (Fig. 6). Substitutions in either the DFD patch or rendering one subunit devoid for ATP binding shuts down substrate reduction activity. We propose a model where cooperative interactions between the DFD patch and the nucleotide relieves the auto-inhibition by the N-terminus of BchL and is a key regulatory step in transient assembly of the DPOR complex (Fig. 6). In the related nitrogenase complex, several crystal structures of the homologous Fe-protein have been solved in complex with a variety of nucleotides in the absence and presence of the MoFe-protein. In these structures, the relative distances between the two nucleotides does not change (Supplemental Fig. 3). Since the flexible N-terminal region is not conserved in the Fe-protein of, it may indicate the presence of a different protective mechanism in the nitrogenase system.

**Figure 6.**
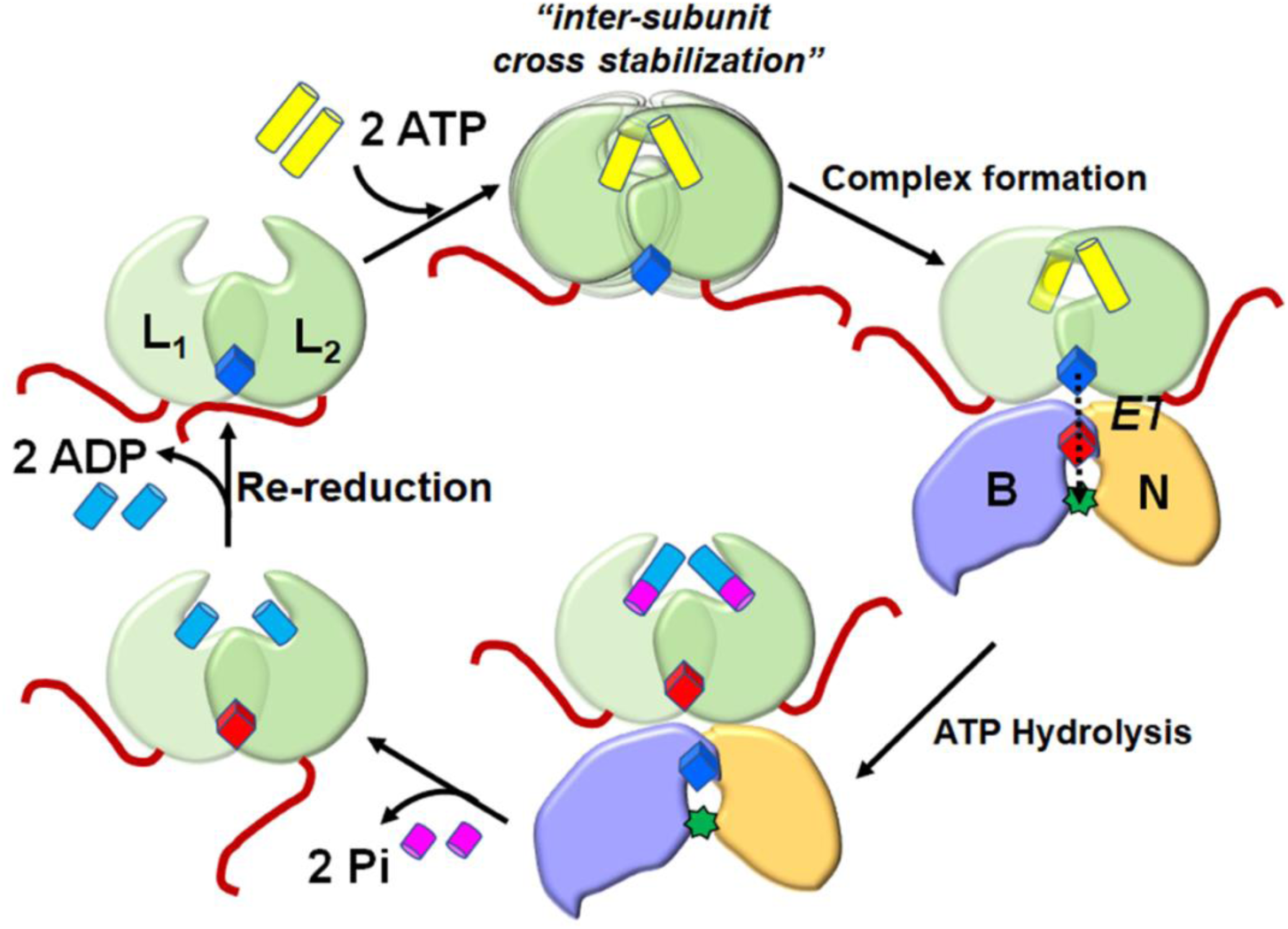
Model for ATP-binding induced release of auto-inhibition by the flexible N-terminus of BchL. Cartoons depict the BchL dimer (green) with one of the two disordered N-termini binding across the [4Fe-4S] cluster (blue; reduced form). ATP (yellow) binding promotes inter-subunit cross stabilization and associated conformational changes through interactions between the DFD patch and ATP; thereby releasing the flexible N-terminus from the docking surface. Complex formation to the BchNB protein ensues followed by electron transfer (ET), ATP hydrolysis, and product (ADP, Pi) release. The oxidized BchL [4Fe-4S] cluster is depicted in red. The BchL [4Fe-4S] cluster is subsequently re-reduced.

In DPOR, the subtle structural changes observed between the ADP-bound and the nucleotide-free crystal structures indicate how nucleotide-binding enables BchL to carry out ET to BchNB. Predictable changes occur around the ATP-binding pocket: Comparison of the two structures suggest that Switch II region may act as a redox switch. The Switch II region is fully conserved between BchL and NifH (the nitrogenase Fe protein), including Phe135 in NifH (Phe163 in BchL), which has been demonstrated to be important for the redox properties of the [4Fe-4S] cluster of NifH.^19^ In the absence of ADP, the local environment immediately surrounding the BchL-cluster is more packed and hydrophobic, promoted by the interactions of Leu155, Val158, and Phe163 (Supplemental Fig. 6). In general, for redox-active metal centers, hydrophobicity increases reduction potential,^20,21^ suggesting that in the absence of Mg-ADP, the BchL-cluster is less likely to become oxidized. While electrochemical data is not available for BchL, studies with NifH show that in the presence of ATP or ADP, the midpoint reduction potential is 120-160 mV more negative,^19^ indicating that oxidation of the [4Fe-4S] cluster of NifH for electron transfer to the P-cluster becomes more favorable in the presence of nucleotides. The conservation of Switch II sequence suggests that the redox properties of the [4Fe-4S] cluster of BchL could be similarly modulated. The van der Waals interaction observed between the [4Fe-4S] cluster of BchL, its ligand Cys160, and Phe163 (Fig. 2c) could be a means of modulating the redox properties of the BchL-[4Fe-4S] cluster in the presence and absence of nucleotides.

Photochemistry and chlorophyll biogenesis occur in the presence of dissolved molecular O_2_ that can generate reactive oxygen species (ROS) such as singlet oxygen and peroxide species. In fact, an increase in BchL transcription occurs when cells encounter singlet oxygen.^22^ The DPOR system thus must maintain functionality in oxygenic environments despite the ability of reactive oxygen species to destroy the [4Fe-4S] cluster necessary for electron transfer. The selective evolution of the flexible N-terminal region and the protective role it plays by binding across the [4Fe-4S] cluster are likely functional adaptations to enable DPOR to function in oxygenic environments. In the case of DPOR, ATP binding relieves autoinhibition. Intriguingly, such protective mechanisms are not found in the Fe-protein of nitrogenase as nitrogen reduction predominantly functions under anaerobic conditions. It is interesting to note that the N-terminal region is not strictly conserved in BchX of the COR enzyme complex, which catalyzes the subsequent reduction of Chlide. Whether the N-terminus of BchX plays a similar role in capping the [4Fe-4S] cluster remains to be established.

## Materials and Methods

### Reagents and Buffers

Chemicals were purchased from Sigma-Millipore Inc. (St. Louis, MO), Research Products International Inc. (Mount Prospect, IL) and Gold Biotechnology Inc. (St. Louis, MO). Oligonucleotides for cloning were purchased from Integrated DNA Technologies (Coralville, IA). Enzymes for molecular biology were purchased from New England Biolabs (Ipswich, MA). All reagents and buffers were thoroughly degassed using alternating cycles of vacuum and nitrogen pressure on a home built Schlenk line apparatus. Anaerobic conditions were maintained via airtight syringes, excess reductant and a vinyl glove box (Coy Laboratories, MI) under a Nitrogen (95%) Hydrogen (5%) mix atmosphere.

### Generation of protein overexpression constructs

The coding regions for BchL, BchN, and BchB were PCR amplified from *Rhodobacter sphaeroides* genomic DNA and cloned into pRSF-Duet 1 or pET-Duet 1 plasmids as described.^12^ Mutations in BchL were generated using Q5 site-directed mutagenesis (New England Biolabs, Ipswich, MA). Plasmids used to express the linked-BchL-proteins carrying glycine linkers of various lengths were synthesized as codon-optimized genes (Genscript Inc., Piscataway, NJ). The longest iteration of the linked-BchL-protein was generated as described in the supplemental section.

### Protein purification

BchL and BchNB proteins were overexpressed and purified as originally described,^23^ with modifications as recently reported.^12^ The following additional steps were added to the purification of the linked-L-proteins. During cell lysis and all subsequent steps, protease inhibitors (protease inhibitor cocktail, Millipore-Sigma Inc – catalog #P2714) and 1 mM PMSF were added to all buffers. As an additional purification step, the concentrated linked-L-protein from the Q-Sepharose eluate was subsequently fractionated over a Sephadex S200 26/600 PG (GE Life Sciences) column using STD buffer (100 mM HEPES, pH 7.5, 150 mM NaCl, 10 mM MgCl_2_, 1.7 mM sodium dithionite, 1 mM PMSF and protease inhibitors). Protein concentrations were determined using the Bradford assay and Bovine Serum Albumin as a standard.

### Generation of Pchlide

Pchlide was generated from a *Rhodobacter sphaeroides* ZY-5 strain harboring a deletion of the BchL gene (a kind gift from Dr. Carl Bauer, Indiana University)^24^ and purified as described.^12^

### Pchlide Reduction Assays

Reduction of Pchlide to Chlide was measured spectroscopically by mixing BchNB (5 µM tetramer), BchL (20 µM dimer), and 35 µM Pchlide, in the absence or presence of ATP (3 mM) in STD buffer with 10 mM MgCl_2_. Substrate reduction experiments were carried out in 200 µl reactions and quenched at the denoted timepoints with 800 µl of 100 % acetone. The acetone/reaction mixture was spun down in a table-top centrifuge at 13,226 x g for 4 minutes. The supernatant was transferred to a cuvette and absorbance scans from 600nm to 725nm were recorded on a Cary 100 UV-Vis spectrophotometer (Agilent Technologies, Santa Clara, CA). Chlide appearance was quantified using the molar extinction coefficient 74,900 M^-1^cm^-1^ at 666 nm. For substrate reduction experiments shown in Figure 3 Chlide appearance was measured in aqueous solution inside a Type 41 macro cuvette with a screw cap (Firefly Scientific, Staten Island, NY). Reactions contained BchNB (1 µM tetramer), BchL (4 µM dimer), 35 µM Pchlide, and ATP (3 mM) in STD buffer containing 10 mM MgCl_2_. Reactions were initiated by addition of degassed ATP via a gas-tight syringe, and spectra were recorded from 400-800nm every 60s as described above. Figure 3B shows absorbance values from difference spectra generated by subtracting timepoints from the first spectra recorded before ATP addition.

### BchL crystallization

BchL crystals were grown anaerobically (100% N_2_ environment with <0.1 ppm O_2_) inside a Unilab Pro glovebox (mBraun, Stratham, NH) at 15 °C using the vapor diffusion method.^25^ All materials and buffers were pre-treated to remove oxygen as previously described.^25^ Initial sparse matrix screens were set up anaerobically using a Mosquito Crystal robotic liquid handler (TTP Labtech, Boston, MA). 1 µL of 200 mM sodium dithionite solution was added to every well to ensure complete removal of any dissolved oxygen. For each drop, 200 nL of well solution and 200 nL of 100 µM dimeric BchL (in 100 mM HEPES pH 7.5, 150 mM NaCl, 10% (v/v) glycerol) were mixed. A crystal was observed after approximately one month with the well solution consisting of 0.6 M sodium chloride, 0.1 M MES:NaOH pH 6.5, 20% (w/v) PEG 4000. Larger volume (3-4 µL) drops of the same well solution and protein concentration in 1:1, 1:2 and 2:1 ratios of protein to well solution yielded large single crystals after ∼2-3 months. Prior to freezing, the well solution was mixed in an equal volume of cryoprotectant solution with a final concentration of 9% sucrose (w/v), 2% glucose (w/v), 8% glycerol (v/v), and 8% ethylene glycol (v/v). Crystals were soaked for a few seconds in the cryoprotectant before being cryo-cooled in liquid nitrogen. A crystal from a drop set up with 2 µL well solution and 2 µL protein solution was used for in-house data collection, while different crystals that grew with 2 µL well solution and 1 µL protein solution were used for synchrotron data collection.

### BchL data collection, processing and refinement

An initial model was built using data collected with an in-house Rigaku MicroMax 007HF X-ray source equipped with a Pilatus 300K detector. A complete dataset at cryogenic temperature (100K) was collected to 2.92-Å resolution, which was integrated using HKL2000^26^ and merged and scaled using SCALA in the CCP4 suite.^27^ Phase determination was initially estimated through molecular replacement (PHASER) using the *Rhodobacter sphaeroides* ADP-bound BchL structure (PDB ID: 3FWY, with all ligands removed) as the search model.^8,28^ A solution was found with two dimers in the asymmetric unit. Following molecular replacement, rigid-body refinement was performed in Phenix.^28^ A starting model was built using AutoBuild and further improved with iterative rounds of model building and refinement using COOT^29^ and Phenix.

Higher resolution data were collected on a different crystal at beamline 17-ID-1 (AMX), National Synchrotron Lightsource-II, at the Brookhaven National Laboratory on a Dectris Eiger 9M detector. Two complete datasets collected at 100K were integrated, scaled and merged to 2.6-Å resolution using HKL2000. The partially refined model from home-source data was used as a molecular replacement model for solving the structure in Phenix. The resulting model was improved through iterative rounds of model building using COOT and Phenix. Data processing and refinement statistics are presented in Supplemental Table 1.

### ATP binding assay

Nitrocellulose membranes, cut into 2 × 2 cm squares, were pretreated with 0.5 N NaOH for 2 min, washed extensively with H_2_O, and equilibrated in binding buffer (100 mM Hepes, pH 7.5, 150 mM NaCl, and 10 mM MgCl_2_). In the reactions (100 μL), BchL (4 μM) was incubated with 1 mM ATP + 0.3 μCi α^32^P-ATP for 10 min at 25 °C, and 20 μL aliquots of the reaction were filtered through the membrane on a single filter holder (VWR Scientific Products). The membranes were washed before and after filtration with 250 μL of nucleotide binding buffer and air dried before overnight exposure onto a PhoshorImaging screen. 1 μL aliquots were spotted onto a separate membrane to measure total nucleotide in the reaction. Radioactivity on the membrane was quantitated on a PhosphorImager (GE Life Sciences). Total ^32^P-ATP bound was calculated using the following equation: [[bound^signal^]/[[total^signal^] x 20]]] x [ATP]

### EPR Spectroscopy

EPR spectra were obtained at 5,10, 17.5, and 30 K on an updated Bruker EMX-AA-TDU/L spectrometer equipped with an ER4112-SHQ resonator (9.48 GHz) and an HP 5350B microwave counter for precise frequency measurement. Temperature was maintained with a Cold-Edge/Bruker Stinger S5-L recirculating helium refrigerator, and an Oxford ESR900 cryostat and MercuryITC temperature controller. Spectra were recorded with either 0.3 G (3 × 10^-5^ T) or 1.2 G (0.12 mT) digital field resolution with equal values for the conversion time and the time constant, 1.0 mW incident microwave power, and 12 G (1.2 mT) magnetic field modulation at 100 kHz. EPR simulations were carried out using Easyspin.^30^

### Samples for EPR Spectroscopy

200 μl EPR samples contained 40 μM BchL (or mutant), 1.7mM dithionite, and, where indicated 3mM ATP, 3mM ADP and/or 20uM BchNB. For a subset of the experiments, EPR experiments were carried out with 20 μM BchNB, 40 μM Pchlide, and, where indicated, 3 mM ATP. Protein samples were prepared and transferred to the EPR tubes in the glove box and stoppered with a butyl rubber stopper. Samples were removed from the glove box and immediately flash frozen in liquid nitrogen and then analyzed by EPR.

## Abbreviations

ET: electron transfer
DPOR: dark-operative protochlorophyllide oxidoreductase
Pchlide: protochlorophyllide
Chlide: chlorophyllide

## Acknowledgements

The authors thank Gabrielle Illava for performing supporting experiments and Phil Jeffrey (Princeton Macromolecular Diffraction Facility) for assistance with crystallographic data collection. Additionally, the authors would like to thank Amanda Byer for her critical reading of the manuscript.

## Author Contributions

EIC, SMSI, MBW, SO, RK, KD, BB and MS performed experiments. EIC, SMSI, MBW, JPB, LCS, NA, BB and EA designed experiments, and performed data analysis. EA, NA, BB, and EIC primarily wrote the manuscript.

## SUPPLEMENTAL INFORMATION

**Supplementary Table 1:** Crystallographic data processing and refinement statistics

| Diffraction Statistics |  |
| --- | --- |
| Space Group | I121 |
| Unit cell (Å) | a = 92.51 b = 100.92 c = 117.79<br>$\alpha = 90.00^\circ$ $\beta = 99.27^\circ$ $\gamma = 90.00^\circ$ |
| Wavelength (Å) | 1.008 |
| Resolution Range (Å) | 42.20 – 2.60 (2.69-2.60) |
| Total observations | 303261 |
| Unique reflections | 32733 (3125)* |
| R<sub>merge</sub> | 0.193(0.908) |
| R<sub>pim</sub> | 0.057(0.413) |
| <I/σ(I)> | 9.2(1.5) |
| CC<sub>1/2</sub> | 0.990 (0.573) |
| Completeness (%) | 99.5 (95.3) |
| Multiplicity | 9.3 (5.1) |
| R<sub>work</sub>/ R<sub>free</sub> | 0.231/0.277 |
| Mean B-factor for all atoms (Å<sup>2</sup>) | 85.6 |
| Anisotropy | 0.076 |
| Total number of atoms refined | 7667 |
| Clashscore | 6 |
| Ramachandran favored/allowed (%) | 96.2/3.9 |
| Ramachandran outliers | 0 |
| RMSD of bond lengths (Å)/angles (°) | 0.002/0.393 |
| Sidechain outliers (%) | 2.4 |

\* Values in parentheses refer to the highest resolution shell

**Supplemental Table 2.**
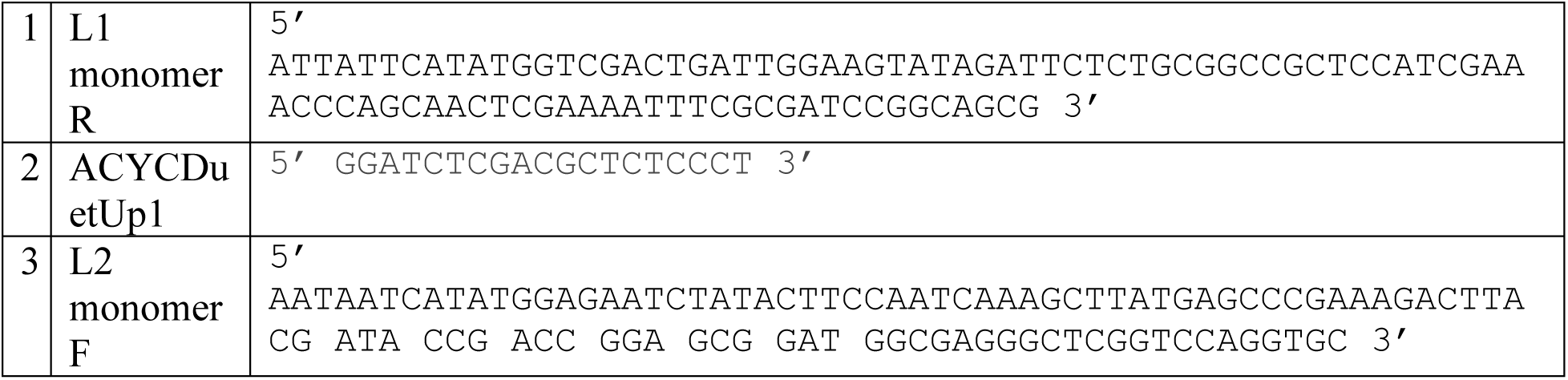
Oligonucleotide primers used in this study

### Supplemental Methods

*Construction of L_1_-L_2_ linked-L-protein constructs:* We used a previously described BchL-RSF-Duet 1 plasmid backbone for cloning the linked-BchL construct.^1^ The two consecutive BchL ORFs are designated L_1_ and L_2_, respectively. For construct 1 – ‘BchL_1_-pRSF-Duet1’, the ORF encoding the L_1_ monomer was PCR amplified using primers #1 and #2 (Supplemental Table 2), designed to eliminate the stop codon at the end of the *BchL* ORF and insert a TEV protease recognition sequence. The PCR product was engineered into an empty RSF-Duet 1 plasmid using BamHI and NdeI. For construct 2 – ‘BchL_2_-pRSF-Duet1’, the ORF encoding the L_2_ monomer was similarly PCR amplified using primer #3 (Supplemental Table 2) and a T7 reverse primer and engineered into an empty pRSF Duet1 plasmid using Nde1 and BamHI. This ORF encodes the linker between the L_1_ and L_2_ genes, as well as a second TEV protease recognition sequence. The flanking TEV sites were designed to remove the intervening linker, post-purification with TEV protease, if necessary. For construct 3 – ‘BchL_1_-L_2_-pRSF-Duet1’, the L_2_ ORF was cut from BchL_2_-pRSF-Duet1 with NdeI and BamHI and engineered into BchL_1_-pRSF-Duet1 using the same restriction sites. Lys44 to Ala substitutions were first introduced into the respective BchL_1_-pRSF-Duet1 or BchL_2_-pRSF-Duet1 plasmids using Q5 mutagenesis and subsequently subcloned into the BchL_1_-L_2_-pRSF-Duet1 plasmid as described above. The subsequent iterations of the linked L constructs (Supplemental Fig. S4c), where the linker lengths were varied, were synthesized as codon-optimized synthetic genes (Genscript Inc.).

**Supplementary Figure 1:**
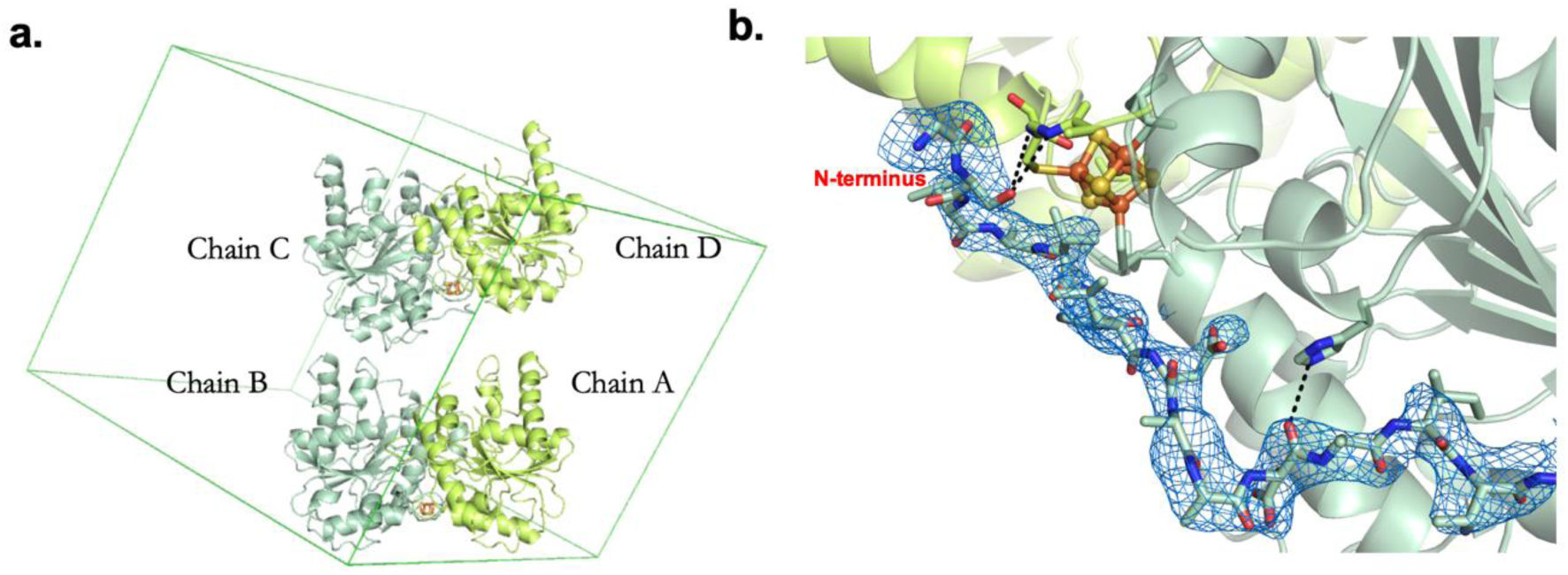
Crystal structure of nucleotide-free BchL **a.** The asymmetric unit of the crystal structure comprises two dimers of BchL chains. The unit cell is outlined in green. **b.** Fo-Fc map of density for the previously unresolved N-terminal tail in chain C (shown as blue mesh, contoured at 1.0σ). Clear, continuous density is visible for the backbone of the entire tail up to residue 16. Interactions of tail residue Ser17 with residues Gly125 and Cys126 on chain D as well as the hydrogen-bonding interaction between Asp26 and His169 of chain C are highlighted.

**Supplementary Figure 2:**
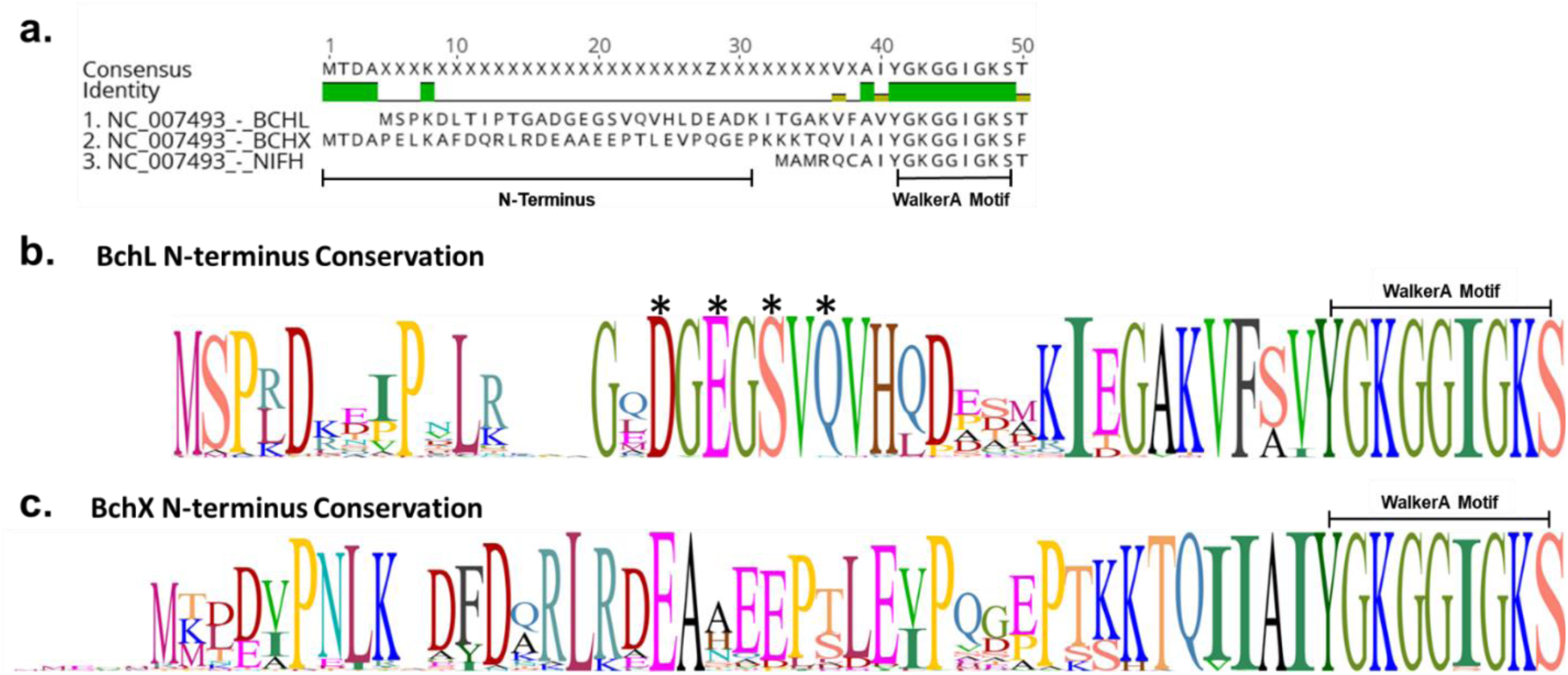
**a.** Sequence alignment BchL, BchX, and NifH (the Nitrogenase Fe-protein). N-terminus length highlighted for BchL and BchX. Walker A motif sequence conservation is also labeled. **b.** Sequence logo of the N-terminus of BchL generated from alignment of n=89 species. Letter height represents sequence conservation. Critical amino acids for capping and generation of BchL^S17A^ and BchL^4A^ are labeled with asterisks. **c.** Sequence logo of the N-terminus of BchX generated from alignment of n=89 species. Letter height represents sequence conservation. Figure demonstrates lack of conservation in N-terminus of BchX. NifH N-terminus conservation is not shown as it lacks enough residues to potentially cap its cluster.

**Supplementary Figure 3:**
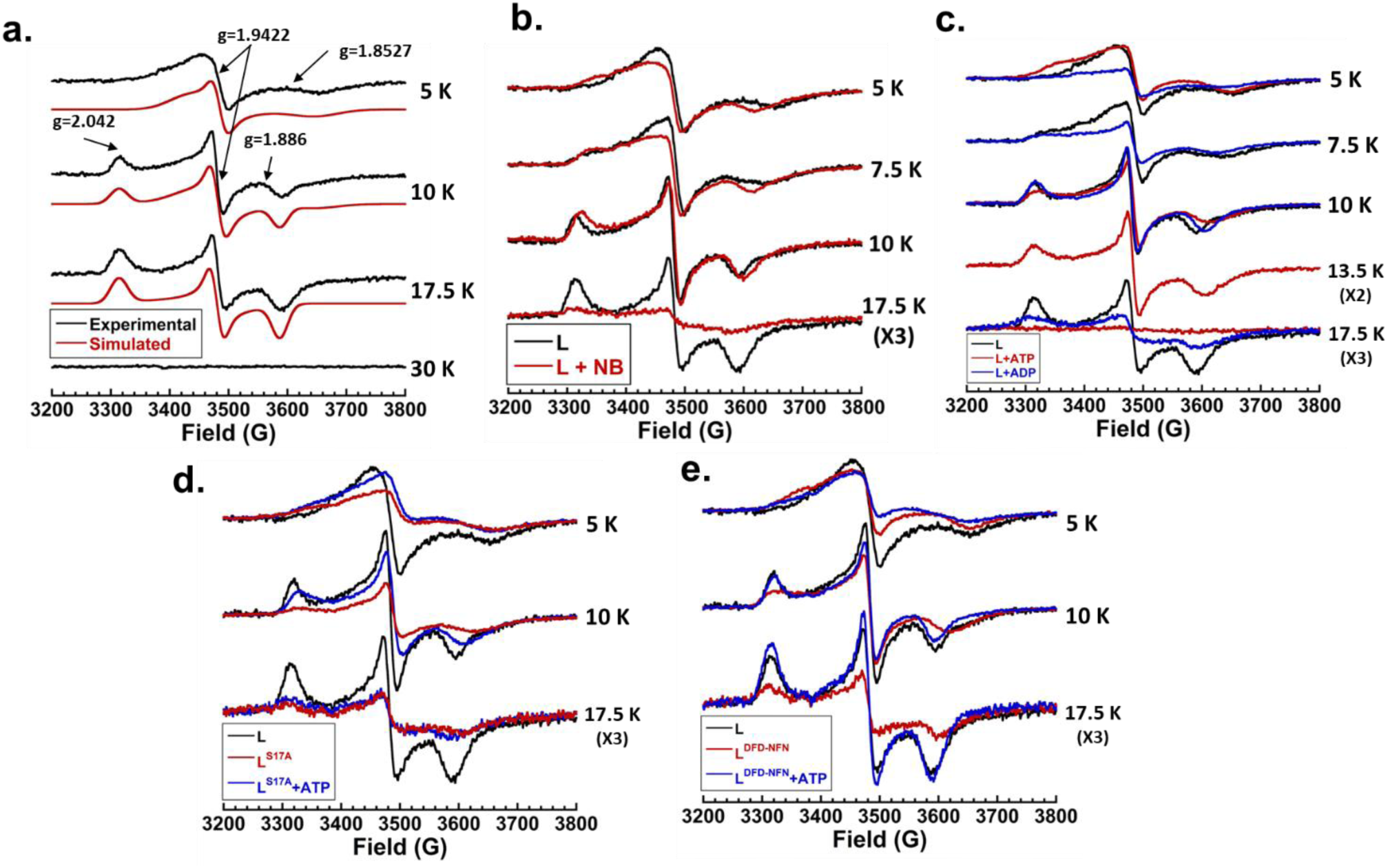
**a.** Experimentally determined (black) and simulated (red) EPR spectra of BchL at 5 K, 10 K, 17.5 K, and 30 K. No absorbance was detected at 35K, typical of [4Fe-4S] clusters. There appear to be two species, a fast-relaxing almost-axial species (g = [1.9750 1.9422 1.8527]) and a slower-relaxing rhombic species (g = [2.0420 1.9442 1.8860]). The spectrum at 10 K was simulated using a 40 % axial:60 % rhombic mixture. The g-values for the axial signal are highly unusual in having all three values less than 2.0. In addition, the line shape could not be replicated well, probably due to rapid passage artifacts. The rhombic signal, on the other hand, is “well-behaved”. **b.** EPR spectra of BchL (black traces) and BchL with 1:1 BchNB (red traces) at indicated temperatures. Upon adding BchNB, there are spectral and relaxation changes. The high-field resonance of the axial signal occurs at a higher g-value (lower field) and features due to the g1 of the rhombic signal persist (distorted by rapid passage) at 5 K. The rhombic signal is very similar in shape to that of the BchL protein alone, but in the presence of BchNB, the signal undergoes significant relaxation broadening at 17.5 K. **c.** EPR spectra of BchL (black traces), BchL incubated with excess ATP (red traces), and BchL incubated with excess ADP (blue traces) at indicated temperatures. ATP incubation produced the most drastic spectral and relaxation changes, particularly at higher temperatures, broadening relaxation almost entirely. ADP incubations produced similar, though not identical effects as BchNB incubation. **d.** EPR spectra of BchL (black traces), BchL^S17A^ (red traces), and BchL^S17A^ incubated with excess ATP (blue races) at indicated temperatures. The slow relaxing axial species was distorted for BchL^S17A^ compared to wildtype. The fast relaxing species was significantly broadened compared to wild-type and was unaffected by ATP incubation, though the mixed-species signal and axial species were both significantly diminished compared to wt. **e.** EPR spectra of BchL (black traces), BchL^DFD-NFN^ (red traces), and BchL^DFD-NFN^ incubated with excess ATP (blue traces). BchL^DFD-NFN^ produced spectra similar to wt BchL with some minor distortions in the axial signal. ATP incubation with BchL^DFD-NFN^ produced spectral changes similar to those produced for the wildtype BchL, though to a lesser extent.

**Supplementary Figure 4:**
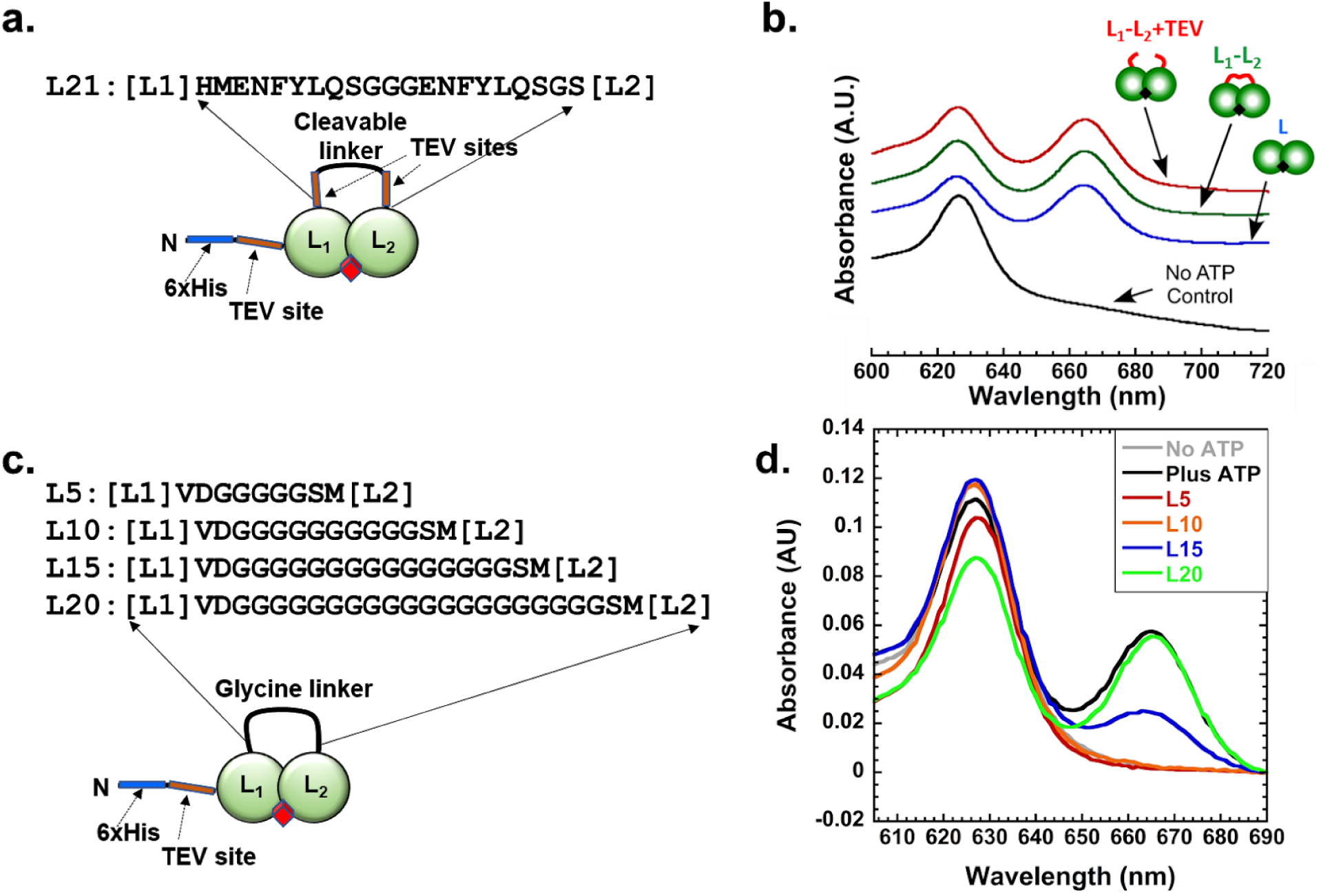
**a.** Cartoon representation of the generated cleavable linked BchL construct, and the amino acid sequence of the linker. **b.** Absorbance plot of acetone extraction of pigments after *in vitro* reaction. Cartoon representations of constructs are shown, and absorbance traces are shifted for clarity. The no ATP (negative control-black), WT reaction (positive control-blue), linked WT (green), and linked WT after cleavage with TEV protease (red) are shown. **c.** Cartoon representation of poly-glycine linkers lacking protease sites, with amino acid sequences of various linker lengths. **d.** Absorbance plot of acetone extraction of pigments after *in vitro* reaction. The no ATP negative control and WT reaction positive control shown as grey and black traces respectively. L5, L10, L15, and L20 construct traces are shown as red, orange, blue, and green lines, respectively.

**Supplementary Figure 5.**
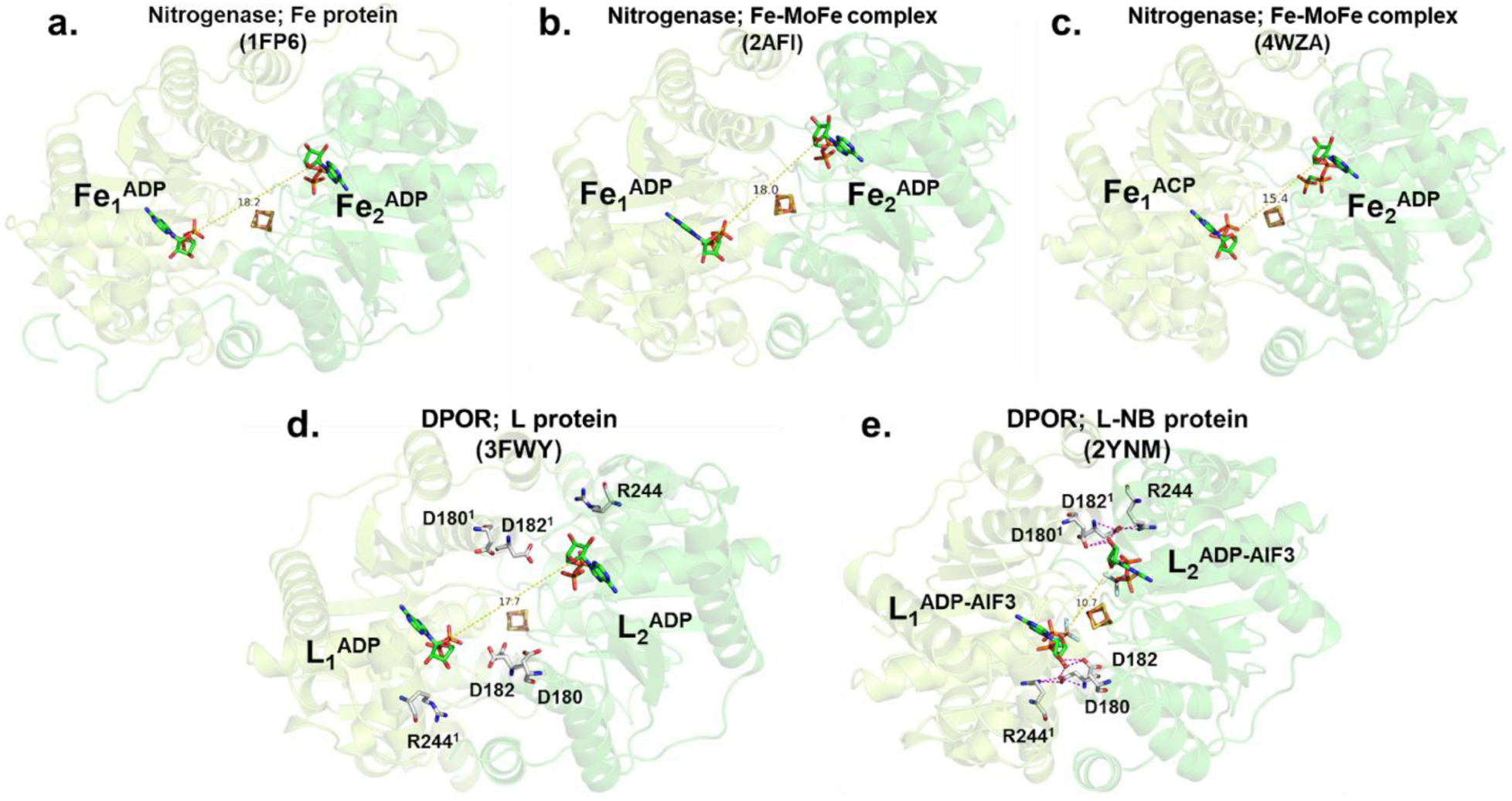
Crystal structures of substrate-bound BchL and the Fe-protein from Nitrogenase. Monomers are colored light and dark green as in previous figures, with highlighted residues and [4Fe-4S] clusters shown as sticks. Distance measurements are shown as dotted yellow lines. **a.** ADP-bound Fe-protein. **b.** ADP bound Fe-protein in complex with Mo-Fe protein**. c.** ADP and ACP bound Fe-protein in complex with MoFe protein. **d.** ADP bound BchL. **e.** ADP-AlF3 bound DPOR in complex with BchNB (not shown).

**Supplementary Figure 6:**
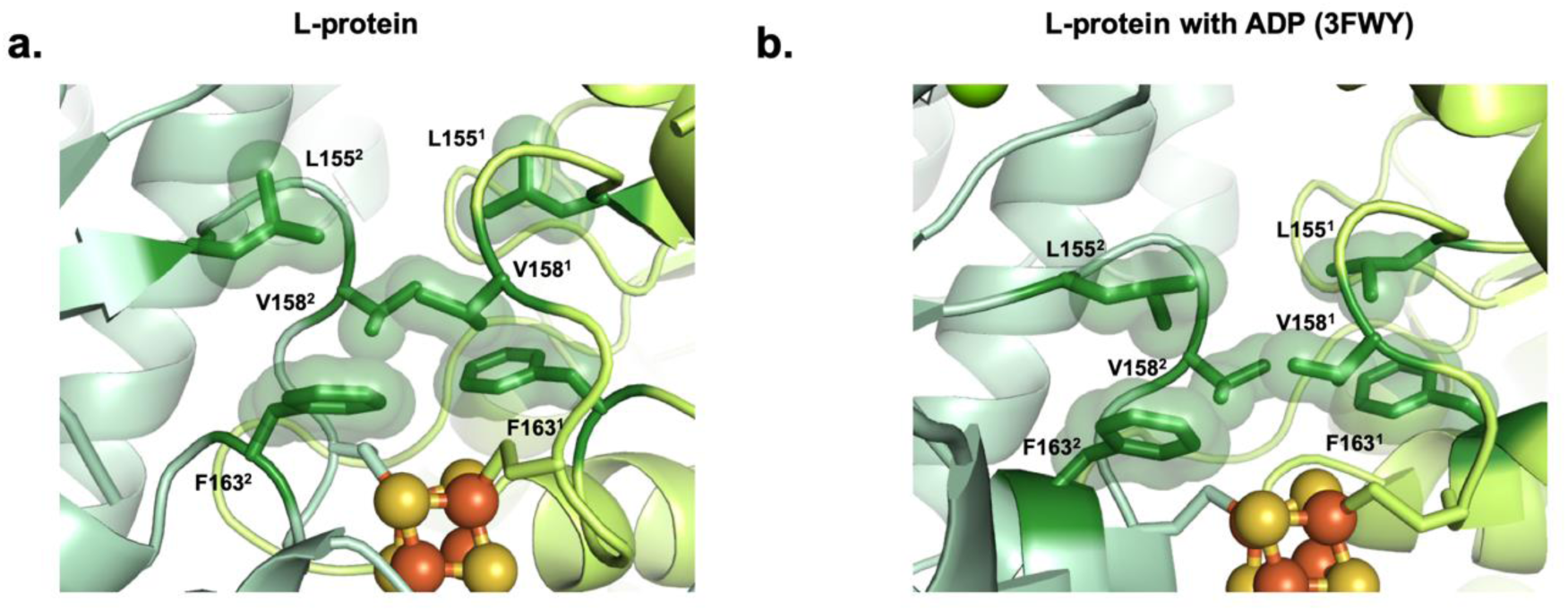
Perturbations in the Switch-II region change the local environment around the [4Fe-4S] cluster. Highlighted in dark green are the interactions of six hydrophobic residues, Leu155, Val158 and Phe163 on each chain, in the absence of nucleotide **a.** and presence of ADP **b.** The environment directly above the [4Fe-4S] cluster becomes less hydrophobic upon the addition of ADP, largely as a result of repositioning of the adjacent Phe163 residues relative to the cluster. More hydrophobic environments generally correlate with increased reduction potential for redox-active metal centers, suggesting that the binding of Mg-ADP acts as a form of redox control in BchL by increasing the tendency of the cluster to become oxidized.

